# Facets of diazotrophy in the OMZ off Peru revisited-what we couldn’t see from a single marker gene approach

**DOI:** 10.1101/558072

**Authors:** Christian Furbo Christiansen, Carolin Regina Löscher

**Affiliations:** Nordcee, Department of Biology, University of Southern Denmark, Campusvej 55, 5230 Odense M, DK; Danish Institute for Advanced Study, University of Southern Denmark, Campusvej 55, 5230 Odense M, DK

## Abstract

Biological dinitrogen (N_2_) fixation is the pathway making the large pool of atmospheric N_2_ available to marine life. Besides direct rate measurements, a common approach to explore the potential for N_2_ fixation in the ocean is a mining based on molecular genetic methods targeting the key functional gene *nifH*, coding for a subunit of the nitrogenase reductase. As novel sequencing and single cell techniques improved, our knowledge on the diversity of marine N_2_ fixers grew exponentially. However, to date one aspect of N_2_ fixation in the ocean is commonly left aside. This is the existence of two alternative types of nitrogenases, which are besides the Mo-Fe nitrogenase *Nif*, the Fe-Fe nitrogenase *Anf*, and the V-Fe nitrogenase *Vnf*, which differ with regard to their metal co-enzymes, as well as regarding their operon structure and composition. *nifH*-based studies may thus be biased, and alternative nitrogenases could not be recovered. Here, we screened a set of 6 metagenomes and -transcriptomes from a sulfidic water patch from the oxygen minimum zone off Peru for genes involved in N_2_ fixation. We identified genes related to all three nitrogenases, and generally increased diversity as compared to our previous *nifH*-based study from the same waters. While we could not confirm gene expression of the alternative nitrogenases from our transcriptomes, we detected additional diazotrophs involved in N_2_ fixation. We suggest that alternative nitrogenases may not be used under conditions present in those waters, however, depending on trace metal limitation in the future they may become active.

**Significance statement:** This study addresses the important process of N_2_ fixation, based on a whole-metagenome and –transcriptome screening, reports an increased diversity of N_2_ fixing microbes in the sulfidic shelf water off Peru, as compared to previous target-gene based studies from the same waters. In addition to a generally higher diversity, genes encoding for alternative nitrogenases, which were previously not subject of any study on N_2_ fixation in oxygen minimum zones, were detected. The ecological meaning and evolutionary history of those alternative nitrogenases is subject of ongoing debates, however, their presence in OMZ waters would allow for N_2_ fixation at extreme anoxia, which may become important in a future ocean challenged by progressive deoxygenation.

## Introduction

Biological fixation of dinitrogen gas (N_2_) is quantitatively the most important external supply of nitrogen (N) to the Ocean. Only certain N_2_ fixing microbes, called diazotrophs, are capable of performing this highly energy costly enzymatic reaction. First pioneering studies involving large scale sequencing surveys, based on the then-available Sanger sequencing technique, identified the paraphyletic nature of diazotrophs throughout the archaeal an bacterial kingdoms (Zani et al. 2000; Zehr, Mellon, and Zani 1998; Zehr and Turner 2001). More recent studies using high throughput sequencing approaches (e.g. (Cheung et al. 2016; Gaby et al. 2018)) broadened the tree of diazotrophs and more clades could be added to the diversity of N_2_ fixers in the Ocean, however, it appears that the initial trees did not fundamentally change with regard to their main cluster structure.

One major reason may be in the nature of the available genetic screening methods, which are mostly based on selectively targeting the *nifH* gene, defined as a key functional marker for the operon encoding for the enzyme dinitrogenase reductase (Yun and Szalay 1984). The operon contains, however, two additional structural genes, *nifD* and *nifK*, altogether the *nif* regulon comprises seven operons (Brill 1980). *NifH* genes are often represented in small numbers in the marine realm, and PCR-based detection requires subsequent amplification steps thus introducing certain biases. Thus, in order to approach the diversity of diazotrophs in the environment, it may be helpful to consider other parts of the *Nif* operon to obtain a more complete picture.

An additional problem regarding the molecular screening for nitrogenases is that the common *Nif*-nitrogenase is only one out of three nitrogenases (Bishop, Jarlenski, and Hetherington 1980; Joerger and Bishop 1988; Kennedy et al. 1991). Two alternative nitrogenases were described, one of which is the *Anf* nitrogenase, which is characterized by an iron-iron (Fe) co-enzyme, the other nitrogenase, *Vnf*, is a vanadium (V)-Fe nitrogenase, while, *Nif* has an Fe-Molybdenum (Mo) cofactor. The difference regarding those metal co factors is of particular interest in anoxic or generally oxygen depleted environments. Mo –in contrast to Fe and V-is only available when at least traces of oxygen are present (Anbar and Knoll 2002; Bertine and Turekian 1973; Collier 1985; Morford and Emerson 1999). This is an important thought with regard to both, the oxygen development in Earth history, and the predicted loss of oxygen in today’s Oceans in a warming world (Keeling, Kortzinger, and Gruber 2010; Schmidtko, Stramma, and Visbeck 2017; Stramma et al. 2008). The question of the evolutionary development of those three nitrogenases is still hotly debated, however, the general conclusion is that they have a common origin (Boyd et al. 2011; Boyd et al. 2015; Boyd and Peters 2013; Raymond et al. 2004), which is supported, e.g. by the structural similarity of at least parts of their operons. Microbes possessing genes for one or both of the alternative nitrogenases always also possess genes for the *Nif*-nitrogenase.

A previous study based on whole genome mining identified a 20 fold increase in diversity when including genes encoding for the alternative nitrogenases *Anf* and *Vnf* (McRose et al. 2017) suggesting the importance of those nitrogenases for obtaining a conclusive picture of the diazotroph community in an environment. Still, information on the presence and distribution of alternative nitrogenases in OMZ waters is to our knowledge not available. In the light of Ocean deoxygenation diazotrophs possessing those nitrogenases may however have and increasingly obtain an advantage in waters with severe oxygen depletion, because Mo may become limiting under anoxia, thus disabling the functionality of the *Nif*-nitrogenase. The oxygen minimum zone (OMZ) off Peru is one of the most prominent examples for expanding and progressing deoxygenation challenging the present ecosystem (Stramma et al. 2010). On the Peruvian shelf, anoxic conditions are severe, with locally occurring sulfidic events promoting a microbial community very different from other parts of the OMZ (Callbeck et al. 2018; Löscher et al. 2015; Schunck et al. 2013). Here, the water column is Fe-rich (Löscher et al. 2014), and anoxic close to the surface, thus sustaining almost euxinic conditions. This environment here thus shares some similarities with ancient sulfidic Oceans (Canfield 1998), thus this area could host a specific diazotroph community, which could shed light on nitrogenase evolution, but also on their future development in terms of progressive oxygen loss from those waters.

Both, by including more than one part of the *Nif* operon, and including the alternative nitrogenases *Anf* and *Vnf* we aimed to obtain novel insights into diazotroph diversity in the Ocean. Since sequencing of full metagenomes and metatranscriptomes is now becoming a standard procedure for exploring biodiversity in the environment, unbiased screening for those both complete operons, and alternative nitrogenases we used this approach to screen for unknown diazotrophs. In this study, we thus re-investigated 6 metagenomes and – transcriptomes from a sulfidic patch in the Peruvian OMZ, and compared the obtained diversity with our previous, Sanger sequencing-based, phylogeny.

## Material and methods

We used and reanalyzed datasets collected on a cruise to the eastern tropical South Pacific in 2009/2010. The cruise was carried out in the framework of the collaborative research center SFB 754 ‘Climate-biogeochemistry interactions in tropical Oceans’ on the German research vessel RV Meteor.

We collected samples as previously described (Löscher et al. 2014; Schunck et al. 2013) on the shelf on station #19, 12°21.88’S, 77°10.00’W, were the water column was anoxic from 20 m down to the sediment (124 m) and hydrogen sulfide (H_2_S) was present in the anoxic zone reaching concentrations up to 5 µmol L^-1^.

Samples for salinity, O_2_ and nutrient analysis, including nitrate, nitrite, ammonia and phosphate (NO_3_^-^, NO_2_^-^, NH_4_^+^ and PO_4_^3-^), respectively were collected from a pump-CTD system. On M77/3 this allowed to combine the classical conductivity-temperature-density sensor measurements with O_2_, fluorescence, turbidity, and acoustic doppler counter profiler measurements, and importantly with continuous water sampling over a water column of maximum 350m depth in high resolution. Seabird sensor O_2_ measurements were calibrated with Winkler method measurements; salinity and nutrients were measured directly after sampling according to Grasshoff et al (1999) using an autoanalyzer. Samples for nucleic acid extraction were prefiltered through 10 µm pore size filters (Whatman Nuclepore Track-Etch) and cells were collected on 0.22 µm pore size filters (Durapore Membrane filters, Millipore) using a vacuum pump, with filtration times not exceeding 20 min. Filters were frozen and stored at −80°C.

Molecular analysis for this study is based on a metagenomic and –transcriptomic dataset originally presented in Schunck et al (2013) with a focus on chemolithoautotrophic lifestyles in sulfidic OMZ waters. Briefly, nucleic acids were purified using the DNA/RNA-Allprep kit (Qiagen, Hildesheim, Germany). In order to maximize the sequencing output for messenger RNA, an isolation kit for prokaryotic mRNA (Epicentre) was used to remove ribosomal RNA (rRNA). Bacterial rRNA was further depleted using the Ambion MicrobExpress kit (Thermo Fisher, Waltham, USA). cDNA was synthesized using the Invitrogen superscript III cDNA synthesis kit with random hexameric primers (Qiagen, Hildesheim, Germany) and samples were subsequently sequenced on a GS-FLX (Roche, Basel, Switzerland) at the Institute for Clinical Microbiology (IKMB) in Kiel, Germany. Four samples were pooled per plate resulting in 1,888,768 (DNA) and 1,560,959 (RNA) sequences with an average length of 392 base pairs, accounting for 757,439,211 and 599,103,110 base pairs of sequence information. Sequence datasets are publicly available from the metagenomics analysis server (MG-RAST) under accession numbers 4460677.3, 4450892.3, 4450891.3, 4460736.3, 4461588.3, 4460676.3, 4452038.3, 4460734.3, 4452039.3, 4452042.3, 4460735.3, 4460734.3 and 4452043.3.

Bioinformatic analysis was performed as described in Schunck et al (2013) using the Meta2Pro annotation pipeline (Desai et al. 2013).

Nitrogenase genes were subsequently searched for in all metagenomes and –transcriptomes. Nitrogenase sequences were subsequently subjected to a BLAST search on the NCBI Genbank database, and ClustalW aligned in Mega 7 (Kumar, Stecher, and Tamura 2016). In order to constrain differences to our previous study of the same samples (Löscher et al. 2014), which were based on *nifH* Sanger sequencing, we constructed Neighbor-joining trees. In order to better constrain which parameters determine the distribution of N_2_ fixers in these sulfidic waters we applied simple correlation analysis and a principal component analysis (PCA, Table S1, Fig. S1).

## Results

In this study we used full metagenomes and –transcriptomes from an environment sharing certain characteristics with an ancient Ocean, in order to explore the functional diversity of N_2_ fixers, and the expression of genes involved in it. We compared the results of this presumably unbiased dataset to previous PCR-based studies targeting the classical functional marker for N_2_ fixation, *nifH*. Sampling took place on the shelf off Peru, in a patch of water which was sulfidic at the time, however, sulfidic conditions were reported from this are in other studies as well, thus this condition could possibly be a more or less permanent feature. Figure 1 (A) shows the bottom water oxygen distribution averaged over around 60 years; the path which was sulfidic during our cruise is visibly oxygen depleted even in this integrated plot. The water column was anoxic from 18 m downwards and hydrogen sulfide (H_2_S) was detected along the vertical profile from 27 m downwards. At the same depth (27 m) nitrate and nitrite were depleted, ammonia was accumulated throughout the anoxic water column (Fig. 1). Iron concentrations were reportedly high reaching concentrations up to about 267 nmol kg^-1^ in the water column, mostly in the bioavailable form of Fe(II) (Schlosser et al. 2018), in line with earlier reports from that same area (Hong and Kester 1986).

**Figure 1:**
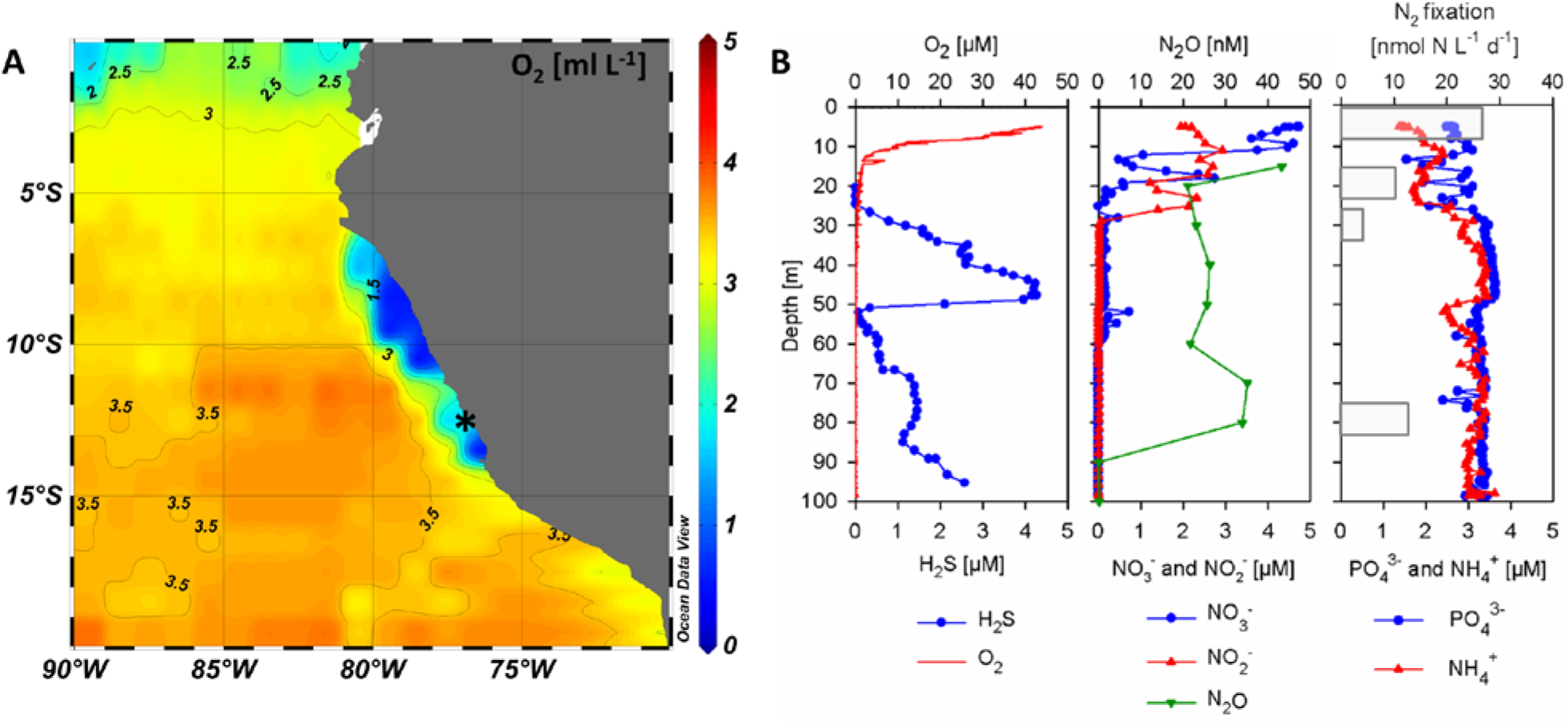
(A) Distribution of bottom water oxygen [ml L^-1^] averaged from 1955-2012 (data from WOA2013). The sulfidic station is indicated with a star. (B) Vertical profiles of O_2_ [µM], H_2_S [µM], N_2_O [nM], NO_3_^-^ [µM], NO_2_^-^ [µM], PO_4_^3-^ [µM], and NH_4_^+^ [µM]. N_2_ fixation rates as presented in Löscher et al (2014) are indicated with grey bars, and were measured using the bubble method (Montoya et al. 1996).

In our previous study from 2014 (Löscher et al. 2014) based on Sanger sequencing of the *nifH* gene, we identified a community of seven previously unknown and two described clades of N_2_ fixers in the OMZ waters off Peru. Organisms matching those clades could be recovered from our metagenomes, thus supporting our previous study regarding the validity of sequence presence (Fig. 2). In addition to those previously described N_2_ fixers, we identified several clades on the genus level from the combined metagenomes and transcriptomes, thus suggesting that the previously used Sanger sequencing did not entirely cover the present diversity.

**Figure 2:**
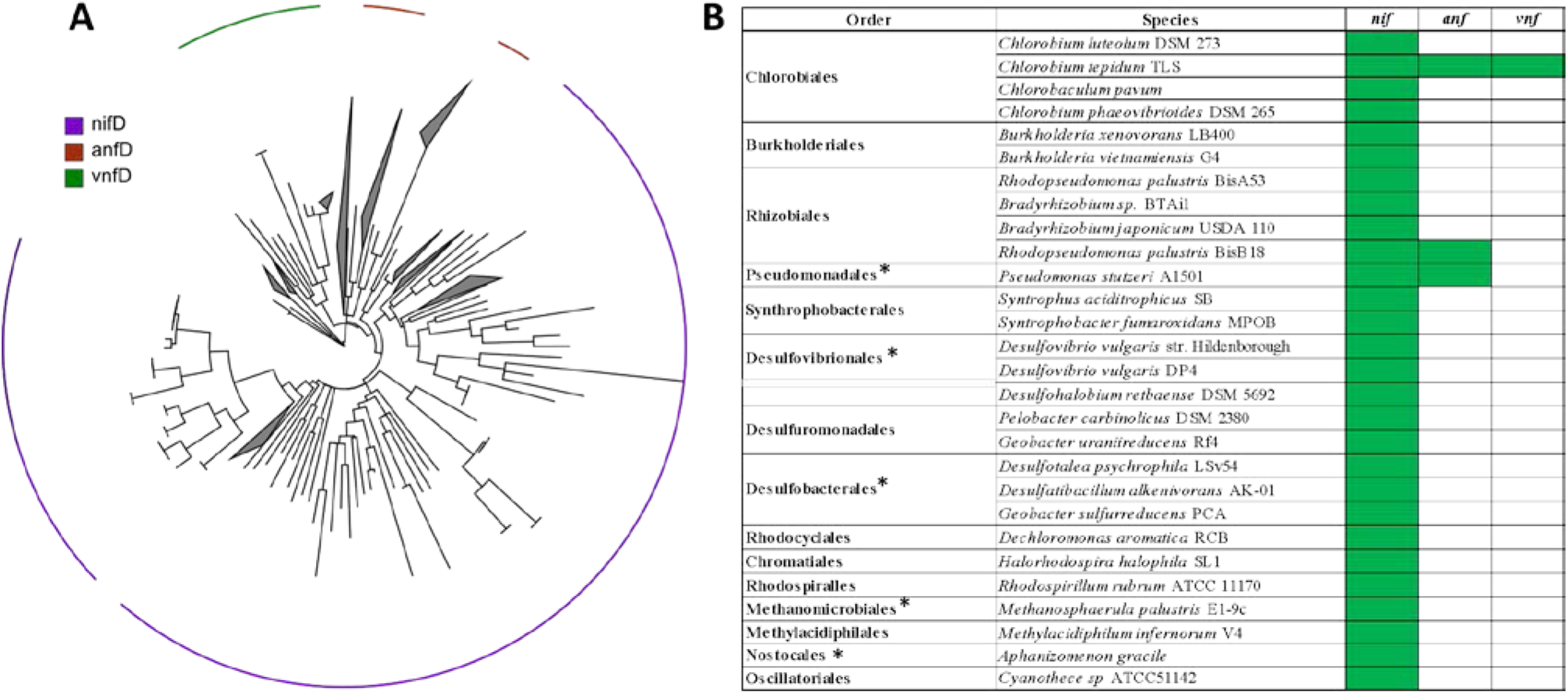
Presence of nitrogenase genes in in the metagenomes and -transcriptomes. (A) The maximum likelihood tree shows the detected diversity based on *nifD*, *anfD*, and *vnfD*. (B) Species closest related to the detected diazotrophs were closest related to the detected diazotrophs, the presence of *nif, anf*, and *vnf* is indicated in green. Note that the majority of diazotrophs possesses *nif*-nitrogenase genes only. * mark genera, of which we found representatives in our previous *nifH*-based study.

Taking all *nif, anf*, and *vnf* genes into consideration, we identified additional N_2_ fixers amongst ß-, ɣ-, and ɗ-Proteobacteria, green and purple sulfur bacteria, Firmicutes, Verrumicrobia, Crenarchaeota and Euryarchaeota. Notably, we report for the first time the presence of alternative nitrogenases in OMZ waters. We detected *anf* genes associated with Pseudomonadales and Rhizobiales, and additional *vnf* genes associated with Chlorobiales, (Fig. 2, 3a).

**Figure 3:**
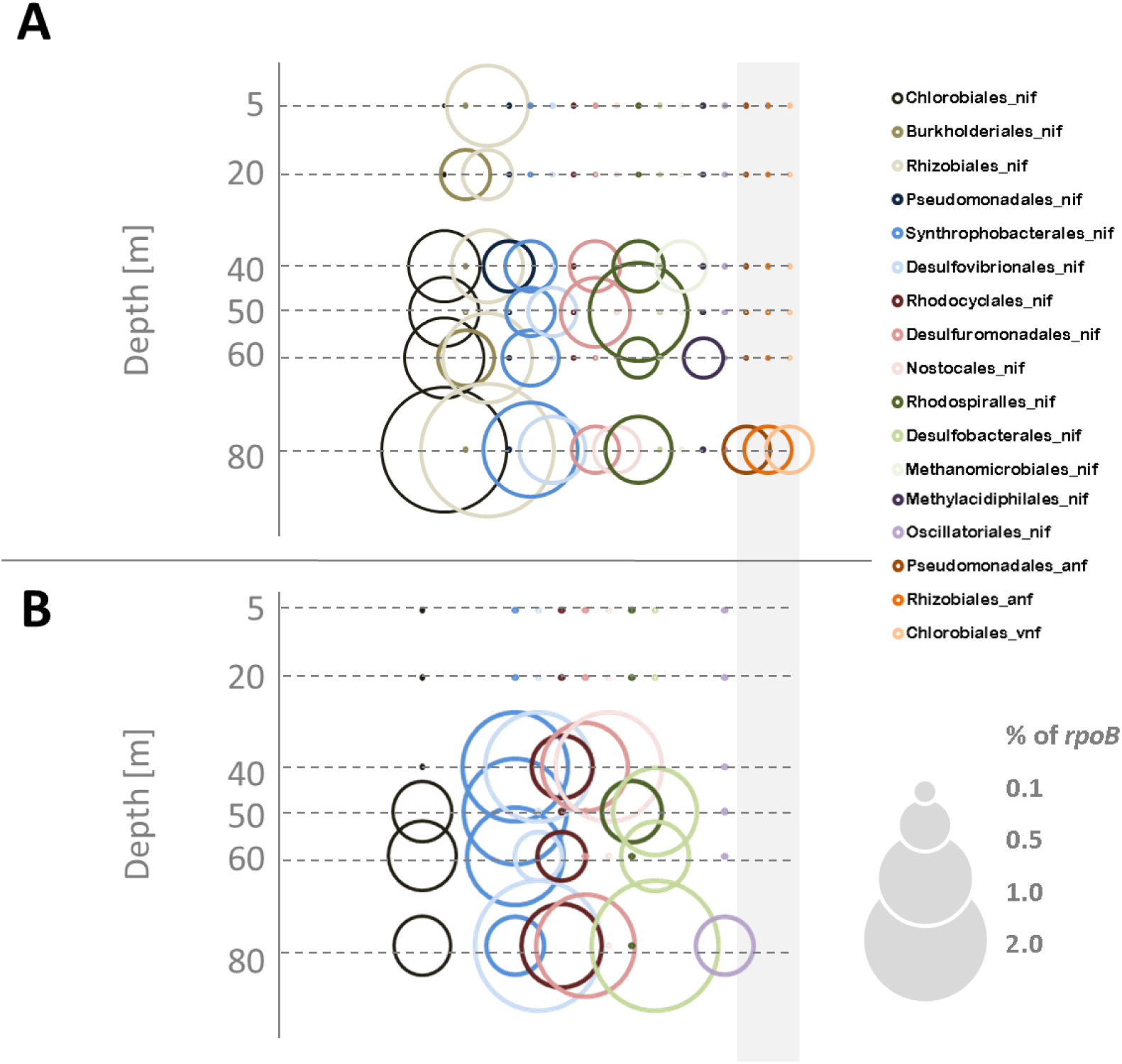
Phylogenetic representation of organisms possessing nitrogenase genes of the *nif*, *anf*, or *vnf* type (all genes of an the operons pooled) in metagenomes and -transcriptomes from the sulfidic station. Genera identified from annotations of protein-coding genes (in the NCBI database) in both our metagenomes (A) and –transcriptomes (B). Nitrogenase gene abundances and expression are shown relative to the putative single copy per organism of RNA polymerase subunit B (*rpoB*). The grey rectangle indicates the alternative nitrogenases *anf* and *vnf*, note that no expression of either of them was found.

Importantly, from none of the identified clades or organisms, we could identify complete operons. This, indeed, leaves us wondering in how far any of the genes identified in this or other studies are translated into functional nitrogenases. On the other hand, N_2_ fixation has been confirmed to occur in those waters at comparably high rates of up to 24.8±8.4 nmol N L^-1^ d^-1^ (Löscher et al. 2016; Löscher et al. 2014). In our previous studies, we could not clearly correlate a specific clade of N_2_ fixers to the high rates in N_2_ fixation in the euphotic zone in the sulfidic patch, however, our data now shows the presence of N_2_ fixers within the genera of both Burkholderiales, Rhizobiales, and Myxococcales in the euphotic zone (Fig. 3a, Table S2), possibly contributing to N_2_ fixation in surface waters.

The metatranscriptomic analysis reveals, however, only an expression of *nif* genes from 40 m downwards, thus again not explaining high surface N_2_ fixation (Fig. 3 b). Identified transcripts only related to *nif*-nitrogenase genes, not to the two alternative nitrogenases. We identified transcripts within the genera of Chlorobiales, Syntrophobacterales, Desulfovibrionales, Rhodocyclales, Desulfuromonadales, and Desulfobacterales in more than one metatranscriptome. The presence and transcriptional activity of the sulfate reducing Desulfovibrionales and Desulfobacterales, as well as the sulfur-respiring Desulfuromonadales, is in line with our previous study, as well as a sediment study on N_2_ fixation from the same area, where those clades have been identified important amongst N_2_ fixers (Gier et al. 2016).

Green sulfur bacteria (Chlorobiales), sulfate-reducing Syntrophobacterales, and Rhodocyclales (mostly classified closest to Dechloromonas clades, which are able to denitrify) have, however, previously not been described to play any role in N_2_ fixation in the Peruvian OMZ. With the exception of the identified Rhodocyclales, all those newly identified N_2_ fixing clades are capable of metabolizing sulfur compounds, thus pointing towards and active sulfur cycle not only in the sediment but also in the water column. However, a statistical correlation to the concentration of H_2_S could only be observed for transcripts of Desulfovibrionales and Desulfuromonadales (Table S2).

Rhodocyclales, in our dataset related to Dechloromonas, are described denitrifiers, which produce the greenhouse gas nitrous oxide as end-product in their denitrification chain (Horn et al. 2005). Transcript abundances were linearly correlated to nitrous oxide concentrations (Table S1), and previous studies reported both, massive denitrification rates, as well as extreme nitrous oxide production from the sulfidic shelf area off Peru (Arévalo-Martínez et al. 2015; Kalvelage et al. 2013).

*Nif* transcripts of Desulfovibrionales and Desulfobacterales were dominant at 80 m depth, which is where N_2_ fixation was at its maximum at anoxic conditions. Those clades were identified as well via *nifH*-based screening, and peaked in abundance at the same depth in our previous study, however, only in the gene, but not in the transcript pool.

## Discussion

By including more genes into our screening for diazotrophs we, expectedly, identified a community more diverse than previously described. We identified clades typical for euxinic environments, such as Chromatiales and Chlorobiales. These clades, besides there potential role for N_2_ fixation are a first evidence for a euxinic state of the Peruvian OMZ. So far, there were two recent descriptions of this area developing sulfidic anoxia (Callbeck et al. 2018; Schunck et al. 2013), however, historical reports of Peruvian fishermen on the characteristic smell and black fishing gear (Schunck et al. 2013), and at least two earlier descriptions of sulfidic anoxia in those waters point towards a re-occurring sulfidic anoxia in this region (Burtt 1852; Dugdale et al. 1977). While the presence of indicator species for euxinia is a curious fact, and their presence in the N_2_ fixer pool may be important in a future with intensifying, expanding OMZs and more frequent events of sulfidic anoxia as predicted for other waters (Lennartz et al. 2014), only the detection of Chlorobium-*nif* transcripts points towards an involvement in N_2_ fixation.

A large fraction, of N_2_ fixers in those sulfidic waters, as previously described (Fernandez, Farias, and Ulloa 2011; Löscher et al. 2014), are microbes involved in sulfur cycling, which appears to be in the nature of this environment, as similarly found in organic carbon-rich sediments (e.g. (Bertics et al. 2013; Gier et al. 2016)). Based on the dominant abundance of transcripts related to Desulfovibrionales and Desulfobacterales, those clades seem to play an important role in N_2_ fixation in those sulfidic waters, possibly contributing to the described high N_2_ fixation rates. In this sense, using full metagenomes /-transcriptomes did, compared to the *nifH*-only based approach, only provide novel insights into identifying active diazotrophs to a certain extent. Tthe power of a metagenome/-transcriptome based approach seems to rather be important of exploring the diversity of possibly underrepresented clades, as well as clades discriminated against by the common *nifH* primers (Zehr, Mellon, and Zani 1998).

One obvious effect of the limitations of a *nifH*-based mining is the previous lack of a description of alternative nitrogenases. In our samples we could identify at least some clades possessing genes for *Anf* and *Vnf*, thus raising the question whether those genes are maintained in the metagenome for at least occasional use or whether they are just leftovers of ancient anoxic situations when Mo was possibly limiting the expression of *Nif*. This, however, can’t be answered from our dataset and is generally a question we can only speculate about. Nevertheless, the differential requirement of trace metals for the three nitrogenases supposedly promotes their expression in dependence of the oxygen status. The turnover of trace metals in general is largely redox sensitive (Morford and Emerson 1999), and has been reported to be impacted by ENSO-dependent changes in redox conditions at the sediment-water interface in this area. With a cyclic alternation between oxic conditions favoring V and Mo fluxes to the water column and anoxic conditions re-precipitating V and Mo to the sediment, both V and Mo accumulate in shelf sediments in the Peruvian OMZ (Scholz et al. 2011). Further, a reduction of V to V(III) by H_2_S has been demonstrated experimentally, thus explaining the redox-dependent accumulation in sulfate reducing sediments (Wanty and Goldhaber 1992). Under sulfidic conditions Mo(VI) is reduced to Mo(IV) and precipitated to the sediments (Crusius et al. 1996). While this precipitation at anoxic and sulfidic conditions in principle removes Mo and V from the water column, it promotes the formation of a reservoir of those trace metals in the sediments, enabling their availability in in the water column via upwelling at least occasionally when redox conditions allow for it. Given the decrease in Mo and V availability at anoxic-sulfidic conditions, possessing the genes for an additional Fe-Fe nitrogenase may thus be advantageous, both regarding ocean anoxic events in Earth history and future ocean deoxygenation likewise. Thus, *Anf* and *Vnf* may presently not play a major role for N_2_ fixation. The detected genetic potential for N_2_ fixation independent of Mo, however, suggests that the Peruvian shelf could sustain its nitrogen demand even under a possible future euxinia.

## Acknowledgements

We acknowledge the captain, crew and the chief scientist, Martin Frank, of the RV Meteor cruise M77/3. We thank the Peruvian government for giving access to their territorial waters. G. Klockgether, A. Ellrott, V. León Fernández, and P. Fritsche, we thank you for technical support during the cruise. The cruise was funded by the German Research Foundation within the Collaborative Research Center SFB754. Funding was further received from the European Union in the framework of the H2020 program (Marie Curie IF, # 704272 to CRL), and from the Villum Foundation (Grant No. 16518; DEC).

